# Validation of cervical lesion proportion measure using a gridded imaging technique to assess cervical pathology in women with genital schistosomiasis

**DOI:** 10.1101/2021.11.16.468781

**Authors:** Katrina Kaestel Aaroe, Louise Thomsen Schmidt Arenholt, Kanutte Norderud, Mads Lumholdt, Bodo Sahondra Randrianasolo, Charles Emile Ramarokoto, Oliva Rabozakandraina, Dorthe Broennum, Hermann Feldmeier, Peter Derek Christian Leutscher

## Abstract

Female genital schistosomiasis (FGS) is characterized by a pattern of lesions which manifest at the cervix and the vagina, such as homogeneous and grainy sandy patches, rubbery papules in addition to neovascularization. A tool for quantification of the lesions is needed to improve FGS research and control programs. Hitherto, no tools are available to quantify clinical pathology at the cervix in a standardized and reproducible manner. This study aimed to develop and validate a cervical lesion proportion (CLP) measure for quantification of cervical pathology in FGS. A digital imaging technique was applied in which a grid containing 424 identical squares was positioned on high resolution digital images from the cervix of 70 women with FGS. A CLP was made for each image by counting the total number of squares containing at least one type of pathognomonic lesions. For validation of inter- and intra-observer reliability, three different observers estimated CLP independently. In addition, a rubbery papule count (RPC) was determined in a similar manner. The intraclass correlation coefficient was 0.94 (excellent) for the CLP inter-rater reliability and 0.90 (good) for intra-rater reliability and the coefficients for the RPC were 0.88 and 0.80 (good), respectively. The CLP facilitated a reliable and reproducible quantification of the surface of the cervix affected by FGS pathognomonic lesions. Grading of cervical pathology by CLP can provide insight into the natural course of schistosome egg-induced pathology of the cervix. Moreover, CLP provides a measure for the efficacy of treatment.

**Author summary:** Female genital schistosomiasis (FGS) is characterized by development of egg-induced chronic inflammatory lesions of the cervix and the vagina. FGS causes various symptoms such vaginal discharge, dyspareunia and post-coital bleeding, and the disease is further associated with reproductive complications such as ectopic pregnancy and infertility. Moreover, FGS is today considered as a major risk factor for transmission of HIV in Sub-Saharan Africa. General prevention directed against *Schistosoma* infection and use of praziquantel as anthelmintic drug therapy are cornerstones in the FGS control strategy. In that overall context, we have developed an important new digital image tool for quantitative assessment of FGS evoked cervical lesions, which enables evaluation of treatment outcome at individual as well as community level. The tool will also provide new information in understanding the natural history of FGS including development of clinical pathology.

## Introduction

*Schistosoma haematobium*, a trematode worm, only occurs in Africa and in the Middle East. In women *S. haematobium* affects particularly the external and internal genital organs. This manifestation is knows as female genital schistosomiasis (FGS).^1^ Millions of women suffer from FGS, which causes a spectrum of genital symptoms signs, including pain during sexual intercourse and post-coital bleeding.^2-4^ Moreover, FGS may lead to infertility and ectopic pregnancy.^5,6^ FGS is associated with a poor quality of sexual life and may cause stigmatization.^7,8^ FGS is also associated with an increased risk of acquiring sexually transmitted pathogens, such as the human immunodeficiency virus (HIV) and the human papilloma virus (HPV).^9-11^

A pocket atlas of the characteristic clinical pathology has been published by World Health Organization (WHO) to assist health-care workers in the diagnosis of FGS in resource-poor endemic areas.^12^ At the cervix, four types of lesions are considered pathognomonic for FGS: grainy sandy patches, yellow homogeneous sandy patches, rubbery papules and neovascularization.^13,14^ According to a consensus recommendation, at least one of the four types of lesions should be present to establish the clinical diagnosis of FGS.^15^

Furthermore, women should undergo colposcopy to ensure proper visualization and identification of the pathognomonic cervical lesions. This examination requires a colposcope, which is an expensive diagnostic instrument requiring maintenance and usually not available in schistosomiasis-endemic areas in rural Africa.^16^ Moreover, colposcopes with an inbuilt digital camera are virtually non-existent. Thus, the interpretation of clinical pathology of the cervix is provided in a descriptive written form at best. Finally, proper use of a colposcope for diagnostic purposes requires extensive training. This makes comparison of changes in cervical pathology difficult. If the clinical pathology of the cervix is not available in a standard digital format, assessment of diagnostic accuracy will be impossible. A digital method to document clinical pathology in women with FGS in a standardized and reproducible manner is imperatively needed. Such a method would be extremely useful in understanding the natural history of the pathology as well as to assess the efficacy of chemotherapy with praziquantel.

We applied a camera equipped with a macro-lens generating high-resolution digital images of the cervix. The images were examined by use of a digital gridded imaging technique to quantify the extension of pathognomonic lesions at the surface of the cervix of women with FGS. The aim of the study was to validate inter- and intra-observer reliability using a cervical lesion proportion (CLP) measure in a series of digital images from women with FGS.

## Methods

The gridded imaging technique study was performed in conjunction with a randomized clinical trial (RCT) in Madagascar assessing the efficacy of different dosages of praziquantel to reduce pathology of the cervix. The trial participants lived in the district of Ambanja in the Northwest region of Madagascar, a highly endemic area for FGS.^17^ In total, 116 participants were recruited from two primary health centers in the municipalities of Antsakoamanondro and Antranokarany. Inclusion criteria for participation in the clinical trial were age 15 to 35 years and presence of FGS pathognomonic cervical lesions. Baseline information on medical history and complaints was obtained from the study participants. Urine samples were collected, and 50 ml of urine were filtrated through a polycarbonate membrane.^18^ After treatment at baseline with praziquantel the study participants were followed up after 5, 10 and 15 weeks involving a clinical re-assessment including photo documentation of the cervical pathology.

### Collection of digital images

A gynecological examination using a speculum was performed at baseline and at each follow-up visit. Photographic documentation of the cervix was obtained using a Canon EOS M50 Camera equipped with a 100 mm macro lens and a circular LED light mounted on the lens. A polarization filter mounted to eliminate reflection of light from the surface of the cervix. The camera was placed on a tripod at 30 cm distance from the surface of the cervix and was mounted on a microscope sledge to allow precise adjustment of the distance. For each patient, the microscope sledge was adjusted to ensure the orifice of the cervix was localized in the center of camera display with a free fringe around the rims of the cervix of 3-5 mm. All digital images captured during the RCT were stored in the REDCap data management system (Research Electronic Data Capture Version REDCap 9.5.6 - © 2020 Vanderbilt University).

### Gridded imaging technique and cervical lesion proportion

The digital part of the study took place in the Centre for Clinical Research at North Denmark Regional Hospital after termination of the RCT. A portfolio of 70 digital images was randomly established from 220 images captured at follow-up at week 5 and 10 among the total 412 images captured in the RCT. Identifiers of the images were anonymized concerning time of image capture to ensure participant confidentiality.

The JavaScript programming software was used to develop a CLP measure based on the quantification method, FGS QubiFier. With the software, a semi-transparent circularly formed grid was superposed as an additional layer to the digital photo of the cervix. The grid covered the whole circumference of the cervix with the orifice appearing in the center of the grid by use of a zoom technique. Thus, the grid covered the total surface of the cervix, thereby enabling to count the grids containing any of the four pathognomonic lesion types in a standardized manner. The grid consists of 424 equally sized squares. Each square in the grid containing a pathognomonic lesion in the digital image, was marked and the different types of lesions were counted automatically (Figure 1). The number of marked squares was then divided by the total number of the 424 squares. The percentage portion constitutes the CLP in the range of zero to 100 as a quantitative estimate of the distribution of lesions covering the cervical surface aiming to identify proper CLP intervals for grading the cervical pathology as mild, moderate, or severe. The number of rubbery papules was also counted and registered as rubbery papule count (RPC).

**Figure 1.**
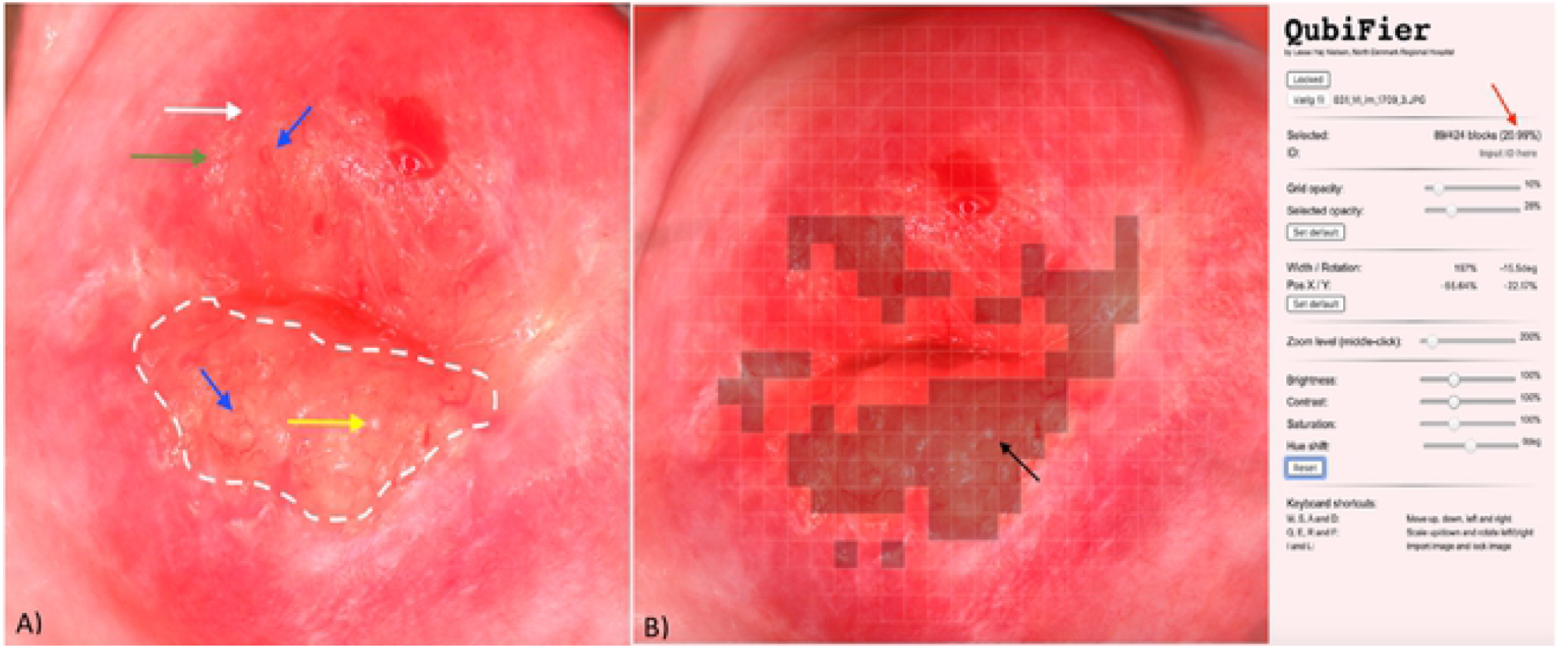
Digital image showing female genital schistosomiasis associated pathognomonic lesions of the cervix (picture A) and the same lesions being digitally marked by QubiFier to determine the cervical lesion proportion (picture B). *The left picture (A) shows an area, which contains homogenous yellow sandy patches appearing as yellow colored area (indicated by white dashed line). The differently colored arrows indicate the other pathognomonic lesion types: a grainy sandy patch (<u>white</u> arrow), clustered grainy sandy patches or rice-grain shaped sandy patches <u>(green</u> arrow) and a rubbery papule colored in beige with an uneven surface <u>(yellow</u> arrow). Abnormal blood vessel (rounded, uneven-calibered, corkscrew or convoluted) are indicated by the blue arrow. The right picture (B) shows squares of the grid marked digitally containing any types of pathognomonic lesions. The red arrow shows the proportion of the cervix covered by any type of pathognomonic lesions.*

### Type of pathognomonic lesions

The following four types of lesions were considered as pathognomonic for FGS^13,19^ and displayed in the WHO FGS pocket atlas.^12^

1. *Grainy sandy patches*, defined as areas with distinct single or clusters of oblong grains (approximately 0.05 × 0.2 mm) in the cervicovaginal mucosa.
2. *Yellow homogenous sandy patches*, defined as homogenous foci without detectable grains at 15 times the original magnification.
3. *Rubbery papules*, previously described in the urinary bladder mucosa only, defined as spheroid, firm, beige, smooth papules (size, 0.3–1.2 mm) in the cervicovaginal mucosa.
4. *Neovascularization*, indicating pathological changes of small venules of corkscrew, unevenly calibrated and convoluted appearance.^20,21^ The distorted blood vessels are the result of eggs released by a female worm which did not achieve to penetrate the wall of the blood vessel and hence become trapped in a venule, thereby blocking the natural blood flow.

### Study design

Three observers (A, B and C), physicians trained and practicing medicine in Denmark, took part in the gridded imaging technique study. The observers had different levels of knowledge and experience regarding gynecology in general and FGS in particular. The two experience levels were graded as minor (= 1), moderate (= 2) and major (= 3), respectively. The experience profiles for the three observers were: A (1 + 1); B (3 + 1); and C (2 + 3).

The three observers analyzed the 70 randomly selected digital images (Figure 2). In an initial consensus rating exercise (Step 1), each of the three observers independently examined ten randomly selected images (images 1 to 10) and rated the cervical lesions. The three observers then shared and discussed their findings to reach consensus on the uniform rating of the images. Subsequently, the remaining 60 images (images 11 to 70) were rated independently by the three observers testing inter-observer reliability (CLP and RPC) in step 2a. Two weeks later intra-observer reliability was tested for images 41 to 70 in a similarly blinded manner by observers A (CLP) and C (RPC) in step 2b. The 60 images used for intra- and interobserver reliability testing were captured from 50 different women: one image from 40 women each (n=40) at either the week 5 or week 10 follow-up visit and two from 10 women at both visits (n=20).

**Figure 2.**
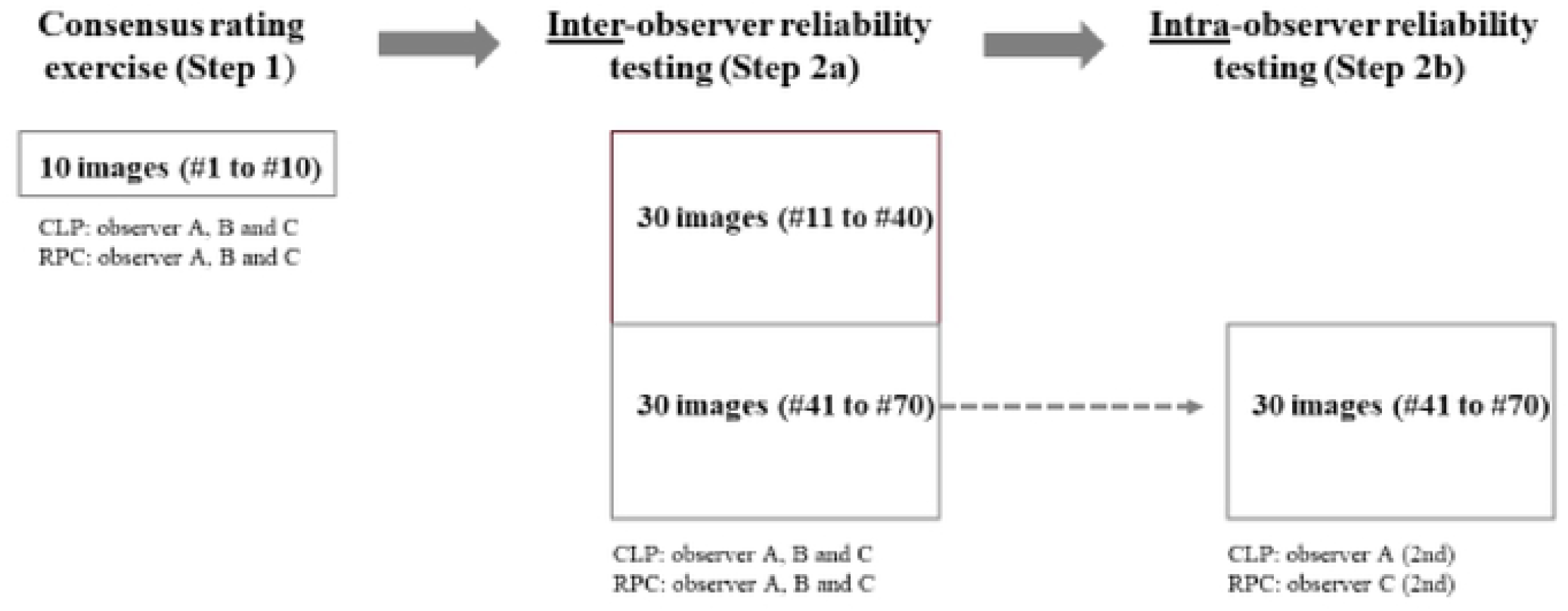
Schematic illustration of the different steps of the study, including the consensus rating exercise and the reliability testing. CLP (cervical lesion proportion); RPC (rubbery papule count)

### Quality of the digital images

Unfortunately, the surface of the cervix could not always be depicted completely for anatomical reasons, e.g. when the labia majora were partially constricted or the cervix was anteverted or retroverted. The impact of the image quality on the CLP rating was therefore assessed between the three observers in separate sessions (A and B, A and C, and B and C, respectively). The quality of the digital images was stratified into whether the entire surface of the cervix was depicted optimally or partially.

### Data management and statistical analysis

The collected data were entered into REDCap data management system. Data validity was ensured by two researchers independently. Data analysis was performed by use of the RStudio programming language (version 1.2.5033: © 2009-2019 RStudio, Inc).

The median of the CLP determined by the three observers was compared using the Wilcoxon rank-sum test. To test if the gridded imaging technique were able to distinguish between clinical pathology including rubbery papules and without rubbery papules, Fleiss kappa was calculated. The value of Fleiss kappa was interpreted as follows: <0.20 (poor), 0.21 to 0.40 (fair), 0.41 to 0.60 (moderate), 0.61 to 0.80 (good) and 0.81 to 1.00 (very good).^22^

The intraclass correlation coefficient (ICC) was calculated by a two-way mixed-effects model based on average rating values and absolute agreement. Intra-observer variation was calculated by a two-way mixed-effects model based on a single-rating and absolute agreement. The ICC values were interpreted as follows: ≤ 0.5 (poor), 0.5 to 0.75 (moderate), 0.76 to 0.9 (good), and > 0.90 (excellent).^23^

### Ethics

The study was carried out in accordance with the Declaration of Helsinki and the Guideline for Good Clinical Practice. Approval was obtained from the Committee of Ethics at the Ministry of Health in Antananarivo (Comité d’Ethique de la Racherche Bio-Médicale auprès du Ministère de la Santé Publique), Madagascar. Informed consent was obtained from all the participating women, including gynecological examination, specimen sampling and image-documentation of clinical pathology.

## Results

### Age and parasitology

The 60 randomly selected images used for intra- and inter-observer reliability testing covered 50 different women: one image from 40 women (n=40) obtained at either follow-up at week 5 or week 10 and two images from 10 women obtained at both follow-up visits (n=20). Median age of the 50 women in the study was 26.5 years (interquartile range (IQR) 20.8-33.0). The median number of *S. haematobium* eggs per 50 ml of urine was 2.5 (IQR 0-62.0). In 13 women, the filtration of 50 ml urine did not reveal eggs.

### Quality of digital images

In 33 (55%) of the 60 digital images used for intra- and interobserver reliability testing, the surface of the cervix was depicted completely, whereas in 27 images (45%) up to 20% of the cervical surface was not visible. However, the observers still found the quality of images to be adequate for CLP and RPC determination.

### Cervical lesion proportion

In Step 2a (inter-observer testing), the CLP ranged from 0.5% to 59.4% in observer A, from 0% to 49.8% in observer B, and from 0.2% to 47.6% in observer C. Figure 3 shows mean CLP by the three observers for each the 60 digital images. The mean proportions were divided into three interval levels defined arbitrarily as low (1 to 15%), intermediary (16 to 30%) and high (>30%) with a distribution as follows: 68% (n=41), 18% (n=11) and 13% (n=8). Three randomly selected cases representing each of the three levels are displayed in Figure 4.

**Figure 3.**
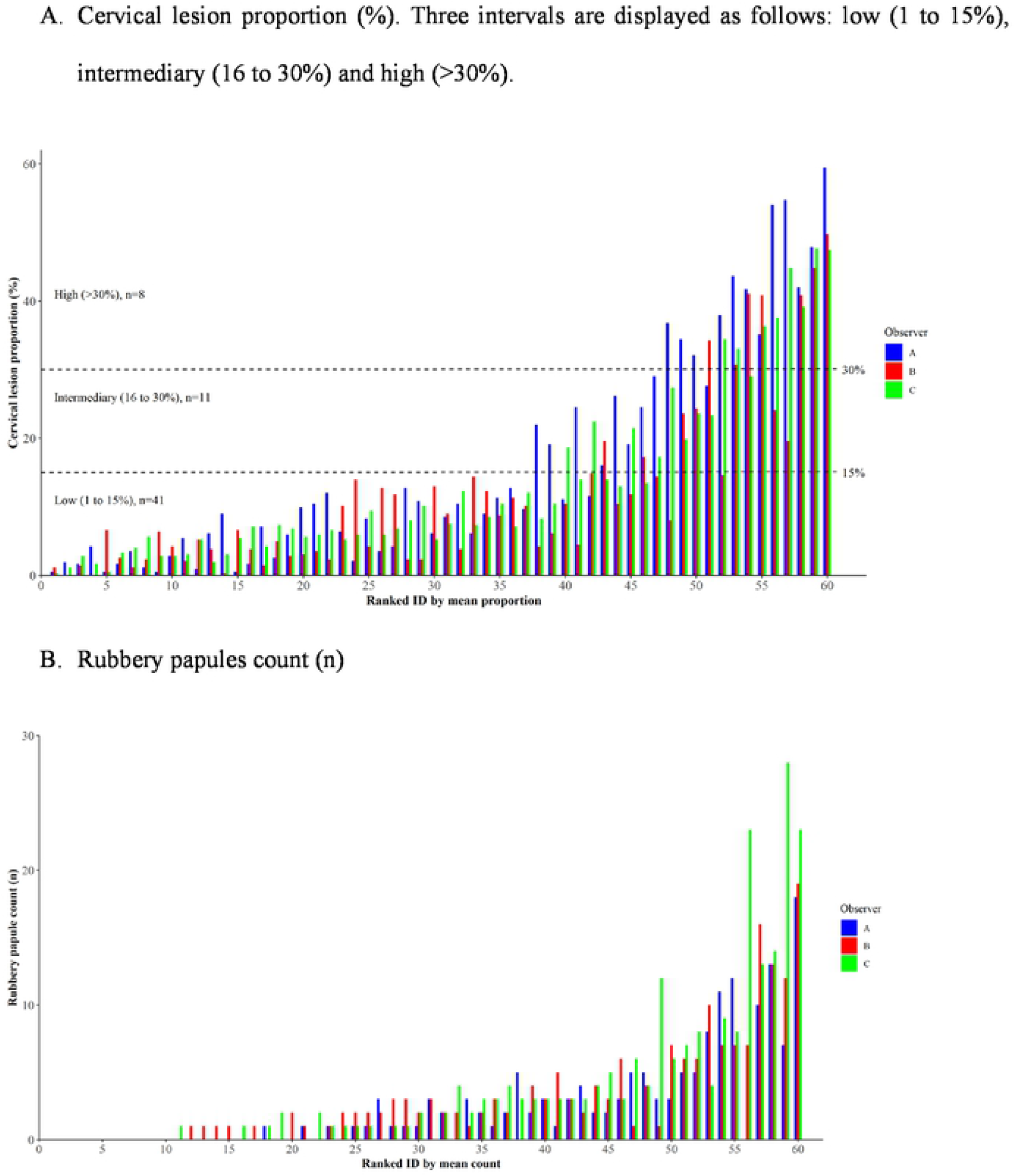
Distribution of cervical lesion proportion (A) and rubbery papule counts (B) for of each the 60 digital images as rated by the three observers and ranked in accordance to the respective individual mean values.

**Figure 4.**
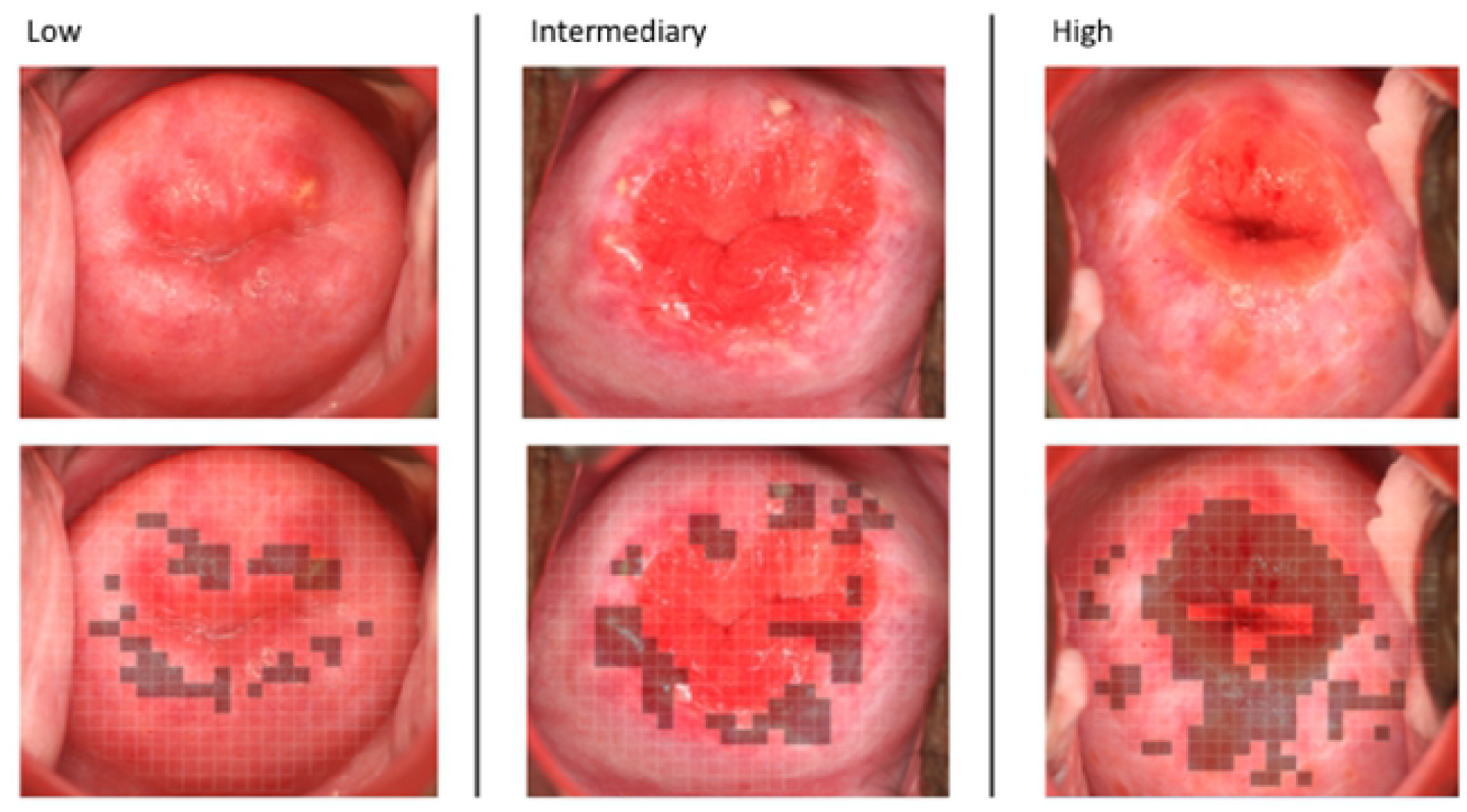
Three cases representing different levels of female genital schistosomiasis (FGS) associated pathology of the cervix in accordance to the cervical lesion proportion (CLP) intervals as follows: low (1 to 15%), intermediary (16 to 30%) and high (>30%).

Median CLP (interquartile range, IQR) for observers A, B and C was 10.1% (3.7-25.8), 8.4% (3.2-14.6) and 7.8% (5.2-19.5), respectively (Figure 5). The difference in scores was statistically significant between observers A and B (p=0.007) and observers A and C (p=0.007), but not between observers B and C (p=0.19). The intra-observer median CLP (IQR) was 6.3% (1.7-11.9) in Step 2a and 8.1% (3.4-14.0) in Step 2b, thus not statistically significant (p=0.352).

**Figure 5.**
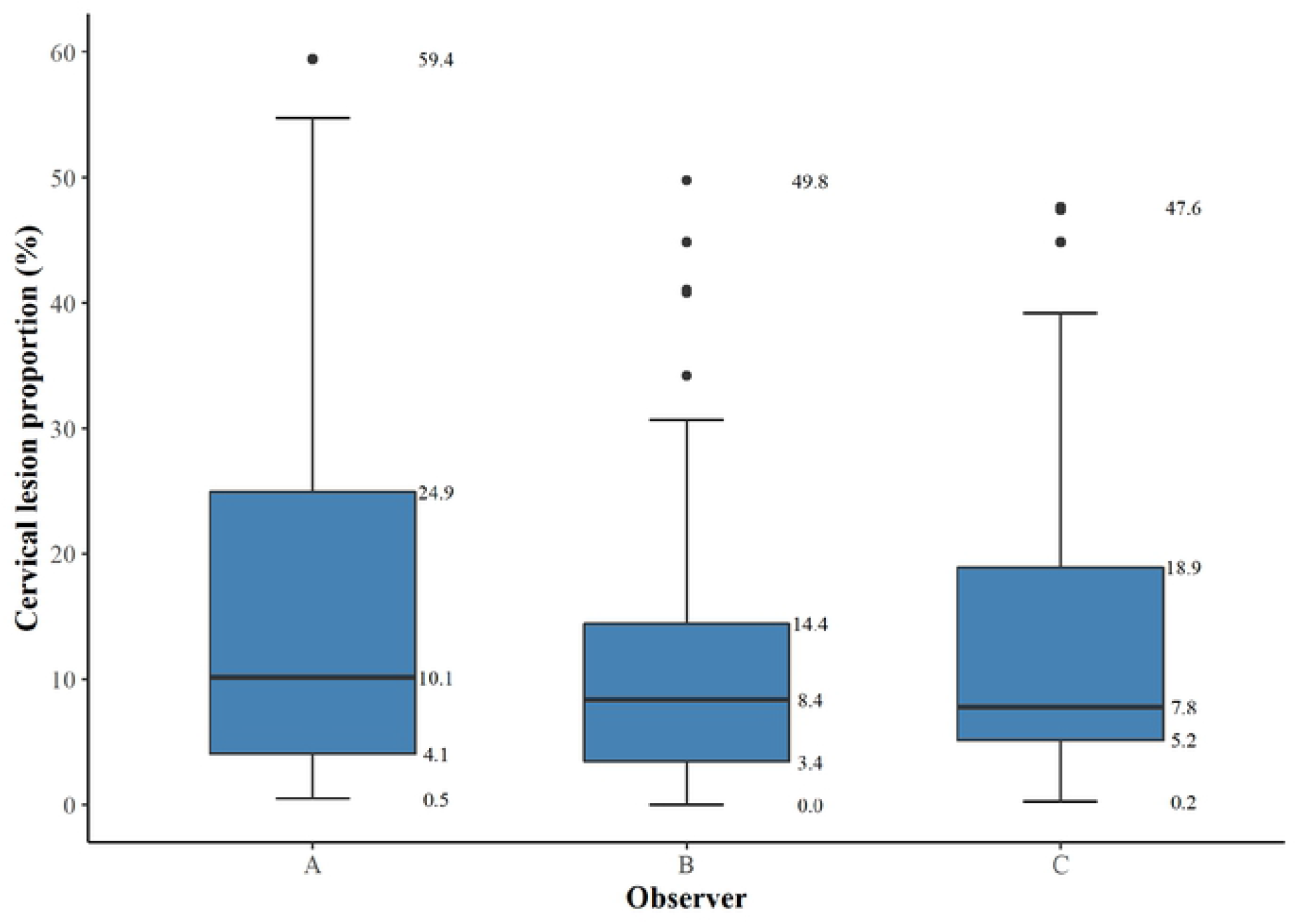
A box-and-whisker plot of cervical lesion proportions (%) for each observer (A, B and C) rating the 60 digital images. The bold horizontal line indicates the median, the upper and the lower line of the box the interquartile range.

### Rubbery papule count

In Step 2a, the RPC ranged from 0 to 18 in observer A, from 0 to 19 in observer B, and from 0 to 28 in observer C (Figure 3). Figure 3 shows the distribution of RPC for each the 60 digital images.

Median RPC (IQR) for observer A, B and C was 1 (0-3), 2 (0-4) and 2 (0-4), respectively (Figure 6). Differences in median scores were statistically significant between observers A and B (p=0.041) and between observers A and C (p=0.024), but not between observers B and C (p=0.438). Median RPC (IQR) was 1 (0-3) in Step 2a and 1 (0-2) in Step 2b, thus statistically significant (p=0.002). The Fleiss kappa value for the three observers in distinguishing rubbery papules was 0.55 (IC 0.54-0.55; p<0001), demonstrating a moderate agreement.

**Figure 6.**
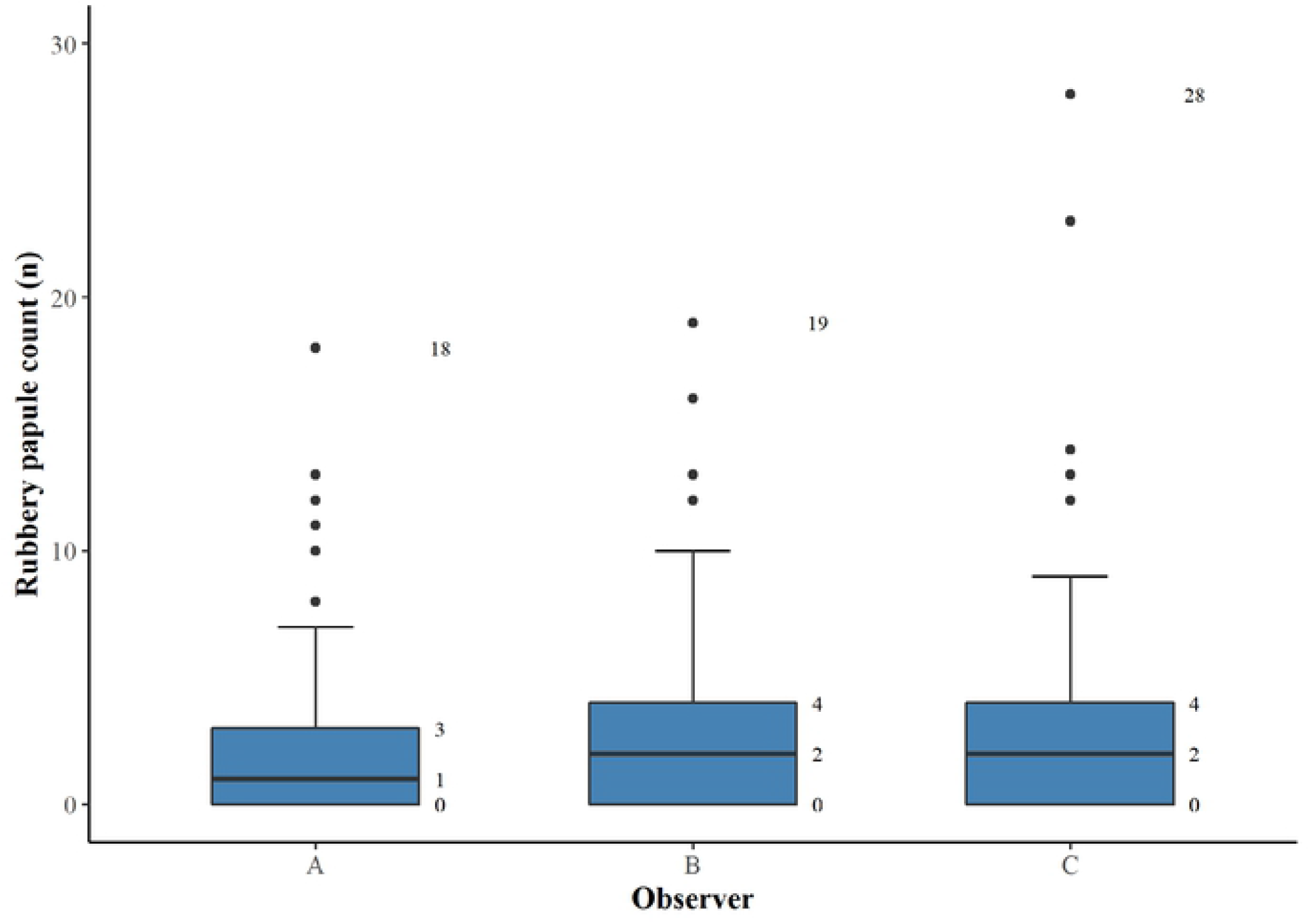
A box-whisker plot of median rubbery papule count (n) and interquartile range for each observer (A, B and C, respectively) rating the 60 digital images.

### Inter- and intra-observer reliability

The inter-observer reliability (A, B and C) for CLP measured by ICC was 0.93 (95% CI 0.90-0.96) indicating an excellent performance. The inter-observer reliability (A, B and C) for RPC measured by ICC was 0.88 (95% CI 0.82-0.92), indicating a good performance. The intra-observer reliability (A) for CLP measured by the intra-class correlation coefficient ICC was 0.90 (95% CI 0.79-0.95), indicating an excellent performance. The intra-observer reliability (C) for RPC measured by ICC was 0.80 (95% CI 0.59-0.90), indicating a good performance.

### Factors influencing discrepancies

Data were further analyzed to identify whether certain characteristics of the surface of the cervix contributed to significant discrepancies in CLP between observers. In ten images with the most pronounced discrepancy in CLP between observers A and B, the differences ranged from 13% to 35%. When comparing observer A with observer C, the differences ranged from 11% to 17% for the ten images representing the most pronounced discrepancy between the two, and for observers B and C, differences ranged between 8% to 25%. Two major domains contributed to discrepancies between observers: 1. difference in zoom-level of the digital images resulting in a different coverage of the surface of the cervix by the grid, 2. disagreement when different cervical lesions co-existed and overlapped in the same square particularly in case of neovascularization. In addition, disagreement occurred also in a few cases due to reduced image quality and/or because the glandular lining of the endocervix was intense and categorized as yellow homogenous sandy patches.

## Discussion

In this study, we developed and validated a method for assessment of FGS pathognomonic cervical lesions based on digital images. Using the gridded imaging technique, we demonstrated that CLP and RPC are measures allowing the quantification of cervical pathology. The method provides an opportunity to further explore the natural history of FGS from a clinical as well as an epidemiological perspective. Moreover, the CLP and RPC offer an opportunity to assess pathological changes following treatment of FGS, in addition to measurement of subsequent re-occurrence of pathology after repeated exposure to *S. haematobium* contaminated water.

The proposed arbitrary grading of cervical pathology as low (1 to 15%), intermediary (16% to 30%) and high (>30 %) according to the CLP, requires further evaluation in other groups of women with FGS. The CLP cut-off points were arbitrarily selected for purpose of the study, and these should be tested clinically t to investigate if different levels of cervical pathology correlate with genital morbidity, including gynecological complaints.

The ICC values for the CLP and RPC indicated good to excellent inter- and intra-observer reliability. The difference in inter-observer reliability concerning CLP (excellent) and RPC (good) can be explained by the difficulty in evaluating a pronounced convex structure such as a rubbery papule on a 2D surface.^14^ A moderate agreement between the three observers was found regarding identification of rubbery papule lesions, emphasizing another important aspect in the validation of the gridded imaging technique. The kappa value demonstrated that the observers were capable of distinguishing between the presence versus absence of this FGS pathognomonic lesion. We believe that rubbery papule lesions represent the sign of recent eosinophil inflammatory response compared to the sandy patch lesions representing a late stage in the natural history of FGS. Thus, a validated RPC could be a good proxy for recent deposition of eggs in the epithelium.

The comparison of CLP and RPC median scores between observers showed no significant difference between observers B and C, whereas a significant difference in median score was seen between observer A and the other two observers. Different levels of experience between the observers could explain this finding, since observer A had less gynecological and clinical FGS experience compared to the other two observers. Two major domains were identified when evaluating the cause of discrepancy between observers. Firstly, different zoom-levels between the observers, and secondly, co-existence of lesions in same grid square. Reduced image quality also played a role in the discrepancy between observers. The possible causes of discrepancies between observers should be addressed in future studies to further optimize the gridded imaging technique protocol.

Use of digital images for the diagnosis and management of FGS provides different advantages in comparison to the colposcopy. FGS pathognomonic lesions are more easily identified on a computer screen as it is possible to enlarge portions of the cervix by increasing the zoom level, thus providing a more detailed visualization of the lesions. Digital images can be revaluated any time and shared with other observers or health care professionals for a second clinical opinion. The digital images of the cervix can be captured by use of the camera alone or by using a camera in conjunction with the colposcopy. A camera with proper technical features may potentially replace the colposcope. In an African context, introduction and use of the colposcope in FGS control programs is challenged by lack of supporting health care and staff training capacity as well as logistic and financial constraints.

Previous studies by Holmen and colleagues have suggested use of colorimetric image analysis as a diagnostic tool in FGS.^16,24^ However, the computer analysis is performed on colposcopic images, which are typically not available in district hospitals in sub-Saharan Africa. The digital camera image documentation approach overcomes the shortage of colposcopic equipment. However, access to cameras may also be limited in rural high endemic areas. Previous studies on visualization of cervical pre-cancer pathology have shown that image documentation could be obtained by mobile phone cameras.^25,26^

### Limitations

This study has some limitations. Firstly, underestimation of the CLS and RPC may have occurred because up to 20% of the cervical surface was visible in half of the images selected for this study. Secondly, colposcopy is currently considered the standard procedure for identification of FGS pathognomonic lesions. In this study, the digital images used for the development and validation of the gridded imaging technique were obtained in women living in an *S. haematobium* endemic area enrolled in a therapeutic trial prior to this study. Main inclusion criteria for participation in this trial were presence of FGS pathognomonic lesions. However, the lesions were identified by digital camera images and not by colposcopy. A combined use of a colposcope and digital camera could have made it possible to study important comparative findings between the two diagnostic methods.

## Conclusion

The gridded imaging technique provides an opportunity to quantify the pathognomonic cervical lesion characteristics in FGS. Furthermore, we found an excellent reliability for the CLP and a good reliability for RPC. Future studies need to explore whether severity of disease as perceived by the patient correlates with CLP and RPC. If so, the gridded imaging technique provides new opportunities in future FGS research and control programs. The study has shown that the WHO FGS pocket atlas constitutes a very useful reference guide for physicians to identify FGS pathognomonic cervical lesions regardless of clinical experience level. Use of WHO FGS atlas together with the gridded imaging technique, allows for potential deployment in *S. haematobium* endemic countries in conjunction with future epidemiological and therapeutic studies aiming at improved FGS control in Africa.

## Acknowledgments

We like to thank the field team for their dedicated work and the women in the district of Ambanja in the Northwest region of Madagascar for willing to participate in the study. Photographer Lasse Høj Nielsen contributed with expertise in development of the gridded clinical image technique,

## Author Contributions

Conceptualization: Louise Thomsen Schmidt Arenholt, Hermann Feldmeier, Peter Derek Christian Leutscher

Data curation: Katrina Kaestel Aaroe, Louise Thomsen Schmidt Arenholt, Kanutte Norderud, Bodo Sahondra Randrianasolo, Charles Emile Ramarokoto, Oliva Rabozakandraina, Dorthe Broennum, Peter Derek Christian Leutscher

Formal analysis: Louise Thomsen Schmidt Arenholt, Mads Lumholdt, Dorthe Broennum, Peter Derek Christian Leutscher

Funding acquisition: Hermann Feldmeier, Peter Derek Christian Leutscher

Methodology: Louise Thomsen Schmidt Arenholt, Mads Lumholdt, Peter Derek Christian Leutscher

Project administration: Bodo Sahondra Randrianasolo, Charles Emile Ramarokoto, Oliva Rabozakandraina, Dorthe Broennum, Peter Derek Christian Leutscher

Resources: Bodo Sahondra Randrianasolo, Charles Emile Ramarokoto, Oliva Rabozakandraina, Peter Derek Christian Leutscher

Software: Louise Thomsen Schmidt Arenholt, Mads Lumholdt, Dorthe Broennum, Hermann Feldmeier, Peter Derek Christian Leutscher

Supervision: Bodo Sahondra Randrianasolo, Dorthe Broennum, Peter Derek Christian Leutscher

Validation: Bodo Sahondra Randrianasolo, Charles Emile Ramarokoto, Oliva Rabozakandraina, Dorthe Broennum

Writing – original draft: Katrina Kaestel Aaroe, Louise Thomsen Schmidt Arenholt, Kanutte Norderud, Peter Derek Christian Leutscher

Writing – review & editing: Louise Thomsen Schmidt Arenholt, Mads Lumholdt, Bodo Sahondra Randrianasolo, Dorthe Broennum, Hermann Feldmeier, Peter Derek Christian Leutscher

